# Transcranial direct current stimulation (tDCS) over the left prefrontal cortex does not affect time-trial self-paced cycling performance: Evidence from oscillatory brain activity and power output

**DOI:** 10.1101/341388

**Authors:** Darías Holgado, Thomas Zandonai, Luis F. Ciria, Mikel Zabala, James Hopker, Daniel Sanabria

**Affiliations:** Department of Physical Education and Sport, Faculty of Sport Sciences, University of Granada, Granada, Spain.; Mind, Brain and Behaviour Research Centre, Department of Experimental Psychology, University of Granada, Granada, Spain.; School of Sport and Exercise Sciences, Endurance Research group, University of Kent, Chatham, UK.

**Keywords:** Endurance Performance, Brain stimulation, time-trial, neuromodulation, cognitive performance

## Abstract

**Objectives:** To test the hypothesis that transcranial direct current stimulation (tDCS) over the left dorsolateral prefrontal cortex (DLPFC) influences performance in a 20-min time-trial self-paced exercise and electroencephalographic (EEG) oscillatory brain activity in a group of trained male cyclists.

**Design:** The study consisted of a pre-registered (https://osf.io/rf95j/), randomised, sham-controlled, single-blind, within-subject design experiment.

**Methods:** 36 trained male cyclists, age 27 (6.8) years, weight 70.1 (9.5) Kg; VO_2max:_ 54 (6.13) ml.min^−1^.kg^−1^, Maximal Power output: 4.77 (0.6) W/kg completed a 20-min time-trial self-paced exercise in three separate sessions, corresponding to three stimulation conditions: anodal, cathodal and sham. tDCS was administered before each test during 20-min at a current intensity of 2.0 mA. The anode electrode was placed over the DLPFC and the cathode in the contralateral shoulder. In each session, power output, heart rate, sRPE and EEG (at baseline and during exercise) was measured.

**Results:** There were no differences (F = 0.31, p > 0.05) in power output between the stimulation conditions: anodal (235 W [95%CI 222 - 249 W]; cathodal (235 W [95%CI 222 - 248 W] and sham (234 W [95%CI 220 - 248 W]. Neither heart rate, sRPE nor EEG activity were affected by tDCS (all Ps > 0.05).

**Conclusion:** tDCS over the left DLFC did not affect self-paced exercise performance in trained cyclists. Moreover, tDCS did not elicit any change on oscillatory brain activity either at baseline or during exercise. Our data suggest that the effects of tDCS on endurance performance should be taken with caution.

## Introduction

Self-paced exercise refers to a physical activity in which the effort needs to be evenly distributed and monitored in order to complete the task without reaching premature exhaustion [1]. Performance in self-paced exercise is undoubtedly related to the functioning of peripheral body systems, such as the muscles, heart, lungs etc., as well as the brain. In this respect, self-pacing during exercise is a challenging cognitive task [2], as it requires constant control and monitoring of internal (e.g., heart rate) and external inputs (e.g., a bump on the road while cycling), while maintaining the goals of the task (e.g. completing a set distance as fast as possible). In other words, self-paced exercise can be regarded as an executive task, with high demands of self-control, goal-monitoring and inhibition [2].

Research in cognitive neuroscience has long pointed to the prefrontal cortex as a key brain area involved in executive processing [3]. Interestingly, the few neuroimaging studies testing participants while exercising have shown activation of the prefrontal cortex, together with the expected sensory-motor recruitment [4,5], which reinforces the hypothesis of the crucial role of executive processing on self-paced exercise. It has been proposed that the prefrontal cortex acts as a control structure by integrating central and peripheral information during exercise, exerting top-down control. The prefrontal cortex would be responsible for merging afferent signals together with inputs provided by the anterior cingulate cortex and the orbitofrontal cortex [6], which has been related to motivational and emotional processing. Therefore, the rationale of the present study was that anodal stimulation of the prefrontal cortex via transcranial direct current would improve self-paced exercise performance, supporting previous evidence (see below).

Transcranial direct current stimulation (tDCS) is a non-invasive electrical brain stimulation technique that is able to induce cortical changes by depolarizing (anodal) or hyperpolarizing (cathodal) a neuron’s resting membrane potential [7]. Recently, there have been an increasing interest in the use of tDCS to enhance endurance performance [8–10]. For example, Angius et al. [9] and Vitor-Costa et al. [10] found an increased time to exhaustion in a cycling test after acute stimulation of the primary motor cortex (M1). Angius et al. [9] attributed that performance enhancement to a reduction of the perceived effort (RPE), although Vitor-Costa et al. [10] did not find such a reduction perceived exertion. These apparently contradictory results leave open the question of whether tDCS affects people’s RPE when stimulating the motor cortex. Meanwhile, Okano et al. [11] found improved cycling performance (greater peak power output) in the anodal condition than in the sham condition after stimulating the temporal cortex of ten trained cyclists. The authors argued that their anodal condition might have influenced activity in the insular cortex, which has been linked to autonomic regulation and to self-perception and awareness of body sensations [12]. Most of research on the effect of tDCS on endurance performance has hitherto been focused on activation or inhibition of the motor and temporal cortices.

To the best of our knowledge, only two studies have targeted the prefrontal cortex. Lattari et al. [13] found increased exercise tolerance in a time to exhaustion at 100% of the peak power after stimulating the left dorsolateral prefrontal cortex for 20-min in eleven physically active women. This improvement was not accompanied by a reduction in the RPE. Meanwhile, Borducchi et al. [14] found an improvement in cognitive performance and mood in elite athletes of different sport modalities (n = 10) after ten days of anodal stimulation over the left dorsolateral prefrontal cortex, which, according the authors, may contribute to performance gains, greater well-being and faster recovery. However, due to the lack of a control condition (Borducchi et al.) and small sample sizes in their studies (like in almost every previous study on tDCS and sport performance), the above results should be considered with caution.

The present (pre-registered, https://osf.io/rf95j/) research is novel as it is the first to directly test the hypothesis that stimulation of the prefrontal cortex would affect performance in a 20-min time-trial self-paced exercise bout in trained male cyclists. More precisely, we expected that activation via anodal stimulation would improve performance, whilst inhibition of the prefrontal cortex via cathodal stimulation would impair performance (compared to a sham condition). The indexes of physical performance were the power output during exercise and the RPE after the self-paced exercise. Additionally, we asked participants to perform an executive task [15] after the exercise. The purpose was to test the hypothesis that any change on physical performance produced by the tDCS over the prefrontal cortex would modulate the subsequent (known [16]) effect of exercise on inhibitory control. This is in line with the idea of a bi-directional relationship between exercise, brain and cognition [16], i.e., brain and cognitive functioning influences exercise performance and vice versa. Brain electrical activity was measured at rest, during exercise, and during the cognitive task by recording electroencephalography (EEG) in order to examine the effects of tDCS at brain level. Even though the literature over the effect of tDCS on EEG is scarce and inconclusive [17], we anticipated an increase in the alpha and beta band after stimulation in the anodal condition compared to cathodal and sham condition.

## Methods

Following institutional ethical approved by the University of Granada Ethics Committee (287/CEIH/2017), a randomized, sham-controlled, single-blind, within-subject experimental design was conducted on male trained cyclists and triathletes with a reported weekly training of more than 7h/week. All experimental procedures were designed to comply with the Declaration of Helsinki.

Before being recruited, participants provided written informed consent having previously read a participant information sheet. All data were entered in a case report form, and subsequently in a computerized database and stored at the Mind, Brain and Behaviour Research Centre (MBBRC) of the University of Granada. Exclusion criteria was the presence of symptomatic cardiomyopathy, metabolic disorders such as obesity (BMI >30) or diabetes, chronic obstructive pulmonary disease, epilepsy, therapy with b-blockers and medications that would alter cardiovascular function, hormonal therapy, smoking, and neurological disorders, as well as the presence of implanted metal devices (e.g., pacemakers, metal plates, wires).

The method and planned analyses of this study were pre-registered on the Open Science Framework. This was done on June 29, 2017, and can be found at https://osf.io/rf95j/.

Additionally, we considered that a medium effect would be appropriate in terms of the potential future practical application of the findings from this type of research to elite cyclists. Therefore, according to the G*Power software [18], 36 participants were required for a power of .8 and a medium effect size, (partial eta-squared *η*^2^ = .13) for a 3 conditions (anodal, cathodal, sham) design. During the data collection, two of the participants could not complete the three experimental sessions and were replaced by two other participants. Accordingly, data collection stopped when complete datasets (successful completion of all three condition) were obtained for 36 endurance trained cyclists and triathletes. The physiological characteristics of the participants are (mean and SD): age = 27 (6.8) years, weight = 70.1 (9.5) Kg; VO_2max_ = 54 (6.13) ml.min^−1^.kg^−1^ and Maximal Power Output: 4.77 (.6) W/kg

Participants visited the MBBRC four times (one screening visit and three experimental sessions). Participants initially attended the MBBRC for a screening visit. After verifying that the participants met the inclusion criteria, they performed a maximal incremental exercise test in order to identify their maximal oxygen consumption using a standard laboratory protocol [19]. After completing the maximal incremental test, participants performed a 10-min time-trial self-paced exercise test in order to familiarise themselves with the protocol to be used in subsequent visits. The shorter duration of the familiarization test (with respect to the proper experimental self-paced exercise) was motivated for the following reasons: 1) our participants were experienced cyclists used to performing self-paced exercise during training and competitions (at high intensity and even for longer durations than that of the experimental self-paced test); 2) most of the participants had already enrolled on previous studies from our lab in which we also used the same test; 3) we were aware that the 10-min test was performed after the maximal incremental exercise test and participants were already fatigued.

After the screening visit, participants attended the lab on three separate occasions to perform the 20-min time-trial acute self-paced exercise (all procedures were the same, except for the stimulation condition). Participants were asked to refrain from drinking alcohol (48 h abstinence) and caffeine (24 h abstinence) and instructed not to perform any exhaustive exercise in the 48 h before each experimental session. Participants were also asked to keep their pre-exercise meal the same for every session. The experimental sessions were completed at the same time of the day to avoid diurnal variations. EEG was recorded throughout the session, except for the stimulation period. Before the beginning of the stimulation, we recorded 5-min EEG with open-eyes as a baseline measure. After the baseline measure, we delivered 20-min of tDCS stimulation: anodal, cathodal or sham. The order of presentation of the three experimental conditions was counterbalanced across participants to control for a potential learning effect. Next, we repeated the 5-min baseline EEG measure with open-eyes. After that, participants performed the 20-min self-paced exercise preceded by 5-min warm-up (at 120 watts) on the cycle ergometer (SRM, Julich, Germany). During the data collection, the SRM broke and we had to replace it for a Phantom 5 ergometer (CyleOps, Madison, USA). The Phantom 5 measure the power output using an on-board power meter PowerTap (PowerTap, Madison, USA) with power accuracy of +/− 1.5%. Every participant completed the time-trial self-paced exercise on the same ergometer: seventeen participants completed the trial on the SRM and nineteen on the Phantom 5. Participants were instructed to achieve the highest average power possible during time-trial self-paced exercise and were freely able to change gearing and cadence throughout. Participants were aware of the elapsed time, but they did not have feedback on performance (wattage and heart rate) during, or after the self-paced exercise. Heart rate was measured continuously throughout the protocol (V800, Polar Electro, Finland). Immediately after exercise, we asked the participant to rate their session RPE (sRPE) [20]. Finally, participants completed a 5-min cool-down and the executive task. The interval between the different sessions was at least 48h to allow the full recovery and to minimize carryover effects.

Stimulation was delivered using battery powered DC stimulators (Newronika S.r.l, Milan, Italy) and delivered through a saline soaked pair of surface sponge electrodes (5 × 5 cm). For the anodal (increased excitability) or cathodal (decreased excitability) we targeted the prefrontal cortex. The anode or cathode electrode was placed over F3 area according to the international EEG 10-20 system [21]. The opposite electrode was placed over the contralateral shoulder area in order to avoid the delivery of current on the participant’s scalp. Current was set at 2 mA and was delivered for 20-min, which has previously been shown to provoke cortical changes [22]. The sham stimulation (control) was similar to the anodal and cathodal stimulation but the device only provided 2mA for 30s after which was turned off without the participant’s awareness. This method replicates the sensory feelings experienced in the tDCS trial (i.e. itching and tingling sensations) and cannot be distinguished from it, whether the stimulation is continued or stopped [23]. The EEG cap was kept over the sponges during stimulation period, but the EEG activity was not recorded. At the end of the session (after completing the cognitive task), participants answered a questionnaire regarding their experience during and after the tDCS sessions [24]. The questionnaire included a set of 19 items (e.g. did you have itching during the stimulation?) scored on a scale that ranged from 0 (no effect at all) to 4 (severe effect).

Participants completed a modified flanker task [15], via use of computer software (E-Prime, Psychology Software Tools, Pittsburgh, PA, USA), to assess inhibitory control, a form of executive processing after the self-paced exercise. Here, the flanker task involves the response to the direction of a central arrow surrounded by other arrows pointing in the same or opposite direction. Congruent trials consist of a central target arrow being flanked by other arrows that faced the same direction (e.g., <<<<< or >>>>>). The incongruent trials consist of the target arrow being flanked by other arrows that faced the opposite directions (e.g., <<><< or >><>>). Participants pressed a button with their left index finger when the target arrow (regardless of condition) faced to the left (e.g., ‘<’) and a button with their right index finger when the target arrow faced to the right (e.g., ‘>’). Each trial started with the presentation of a cross (fixation point) that remained on a steady until the appearance of the target arrows 2 seconds later. The target was presented in the middle of the screen for 150 ms and a response window of 1350 ms was allowed. The next trial started 1500 ms after the response. Total task duration was approximately 7-min. Participants completed one block of 160 trials with equal probability for congruent and incongruent trials, randomized across task conditions. A brief familiarization of the task was included in the screening visit. RT (in ms) and response accuracy (percentage of correct responses) for each stimulus were recorded.

EEG were recorded at 1000 Hz using a 30-channel actiCHamp System (Brain Products GmbH, Munich, Germany) with active electrodes positioned according to the 10-20 EEG International System and referenced to the Cz electrode. The cap was adapted to the individual head size for each participant (mean of 57 cm), and each electrode was filled with Signa Electro-Gel (Parker Laboratories, Fairfield, NJ) to optimize signal transduction. Participants were instructed to avoid body movements as much as possible, and to keep their gaze on the centre of a computer screen during the measurement. Electrode impedances were kept below 10 kΩ. EEG pre-processing was conducted using custom Matlab scripts and the EEGLAB and Fieldtrip Matlab toolboxes. Each period and stimuli for the analysis were detected by triggers sent through a parallel port from the E-prime software to the EEG recorder. EEG data were resampled at 500 Hz, with a butter filter design and bandpass filtered offline from 1 and 40 Hz to remove signal drifts and line noise, and re-referenced to a common average reference. Horizontal electrooculograms (EOG) were recorded by bipolar external electrodes for the offline detection of ocular artefacts. Independent component analysis was used to detect and remove EEG components reflecting eye blinks. The potential influence of electromyography activity in the EEG signal was minimized by using the available EEGLAB routines [25]. Independent component analysis was used to detect and remove EEG components reflecting eye blinks [26]. Abnormal spectra epochs which spectral power deviated from the mean by +/− 50 dB in the 0-2 Hz frequency window (useful for catching eye movements) and by +25 or −100 dB in the 20-40 Hz frequency window (useful for detecting muscle activity) were rejected. On average, 2.25 % of epochs per participant were discarded.

All analyses were completed using statistical nonparametric permutation tests with a Monte Carlo approach. These tests do not make any assumption of the underlying data distribution, are unbiased, and as efficient and powerful as parametric statistics. When statistical significance (p < 0.05) was found, values were corrected by the false discovery rate method. The effect of experimental condition (anodal, cathodal, sham) on self-paced exercise power output, heart rate and RPE were analysed using a within-subject design condition.

Spectral power was analysed using a within-participants’ design with the factor of stimulation (anodal, cathodal, sham). Each period (Baseline, Warming Up, Exercise, Cooling Down) was tested separately for significance. In the absence of strong a priori hypotheses over the frequency range and channels which tDCS may induce a change, we use a stepwise, cluster-based, non-parametric permutation test [27]. The spectral decomposition of each epoch (1s) was computed using Fast Fourier Transformation (FFT) applying a symmetric Hamming window (0.5s) and the obtained power values were averaged across experimental periods.

For the cognitive task, we analysed the event-related spectral perturbation main effects of stimulation (anodal, cathodal, sham) for each stimulus (congruent, incongruent) by applying the cluster-based approach [28]. In order to reduce the possibility that the type II error rate was inflated by multiple comparisons correction, we set an a priori criteria of collapsing data into four frequency bands: Theta (4–8 Hz), Alpha (8–14 Hz), lower Beta (14–20 Hz) and upper Beta 1 (20–40 Hz). Task-evoked spectral EEG activity was assessed by computing event-related spectral perturbation in epochs extending from −500 ms to 500 ms time-locked to stimulus onset for frequencies between 4 and 40 Hz. Spectral decomposition was performed using sinusoidal wavelets with 3 cycles at the lowest frequency and increasing by a factor of 0.8 with increasing frequency. Power values were normalized with respect to a −300 ms to 0 ms pre-stimulus baseline and transformed into the decibel scale [29].

## Results

### Side effects

The intervention was well tolerated and participants reported common side effects such as tingling (anodal: 22%, cathodal: 8% and sham: 11%), or “itchy sensation in the scalp (anodal: 30%, cathodal: 8% and sham: 16%).

#### Exercise performance

The average power output during the time trial self-paced exercise was not significantly different (F(2,34) = 0.31, p > 0.05) between conditions (see Fig 1): Anodal (234 W [95%CI 222 - 249 W]; Cathodal (235 W [95%CI 222 - 248 W] and Sham (234 W [95%CI 220 - 248 W].

**Fig 1.**
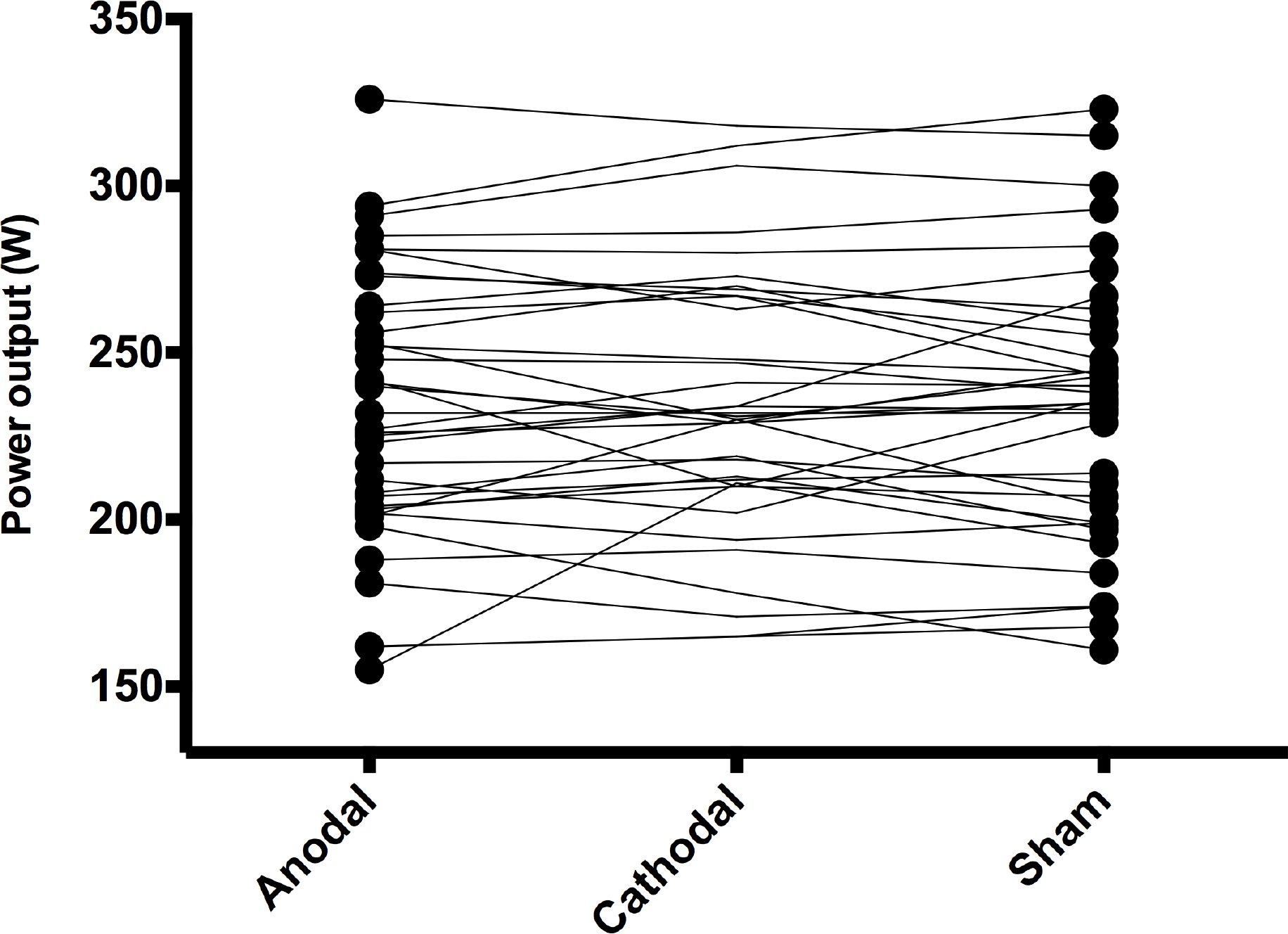
Power output (watts) profile for each participant during the 20-min self-paced exercise.

The heart rate signal for three participants was lost during the 20-min time-trial self-paced exercise, consequently they were removed from the subsequent analysis (n= 33). The average heart rate during the time trial was not significantly different (F(2,34) = 1.02, p > 0.05) between conditions: Anodal (161 beats min−1 [95%CI 157 - 166 beats min−1]; Cathodal (162 beats min−1 [95%CI 158 - 167 beats min^−1^] and Sham (162 beats min^−1^ [95%CI 157 - 167 beats min^−1^].

Post time-trial sRPE did not show any significant differences between conditions: Anodal (17.02 [95%CI 16.5 - 17.5]; Cathodal (17 [95%CI 16.8 - 17.4] and Sham (17.02 [95%CI 16.5 - 17.5], F(2,34) = 1.69; p > 0.05.

#### Electrical brain activity (EEG)

Due to excessive noise in the EEG signal, five participants were not included in the EEG analysis (n= 31). The analysis of tonic spectral power (see Fig. 2) did not provide any significant difference (all ps > 0.05) between conditions (anodal, cathodal and sham), and for each period of time (baseline-pre; baseline-post, warm-up, self-paced exercise and recovery).

**Fig 2.**
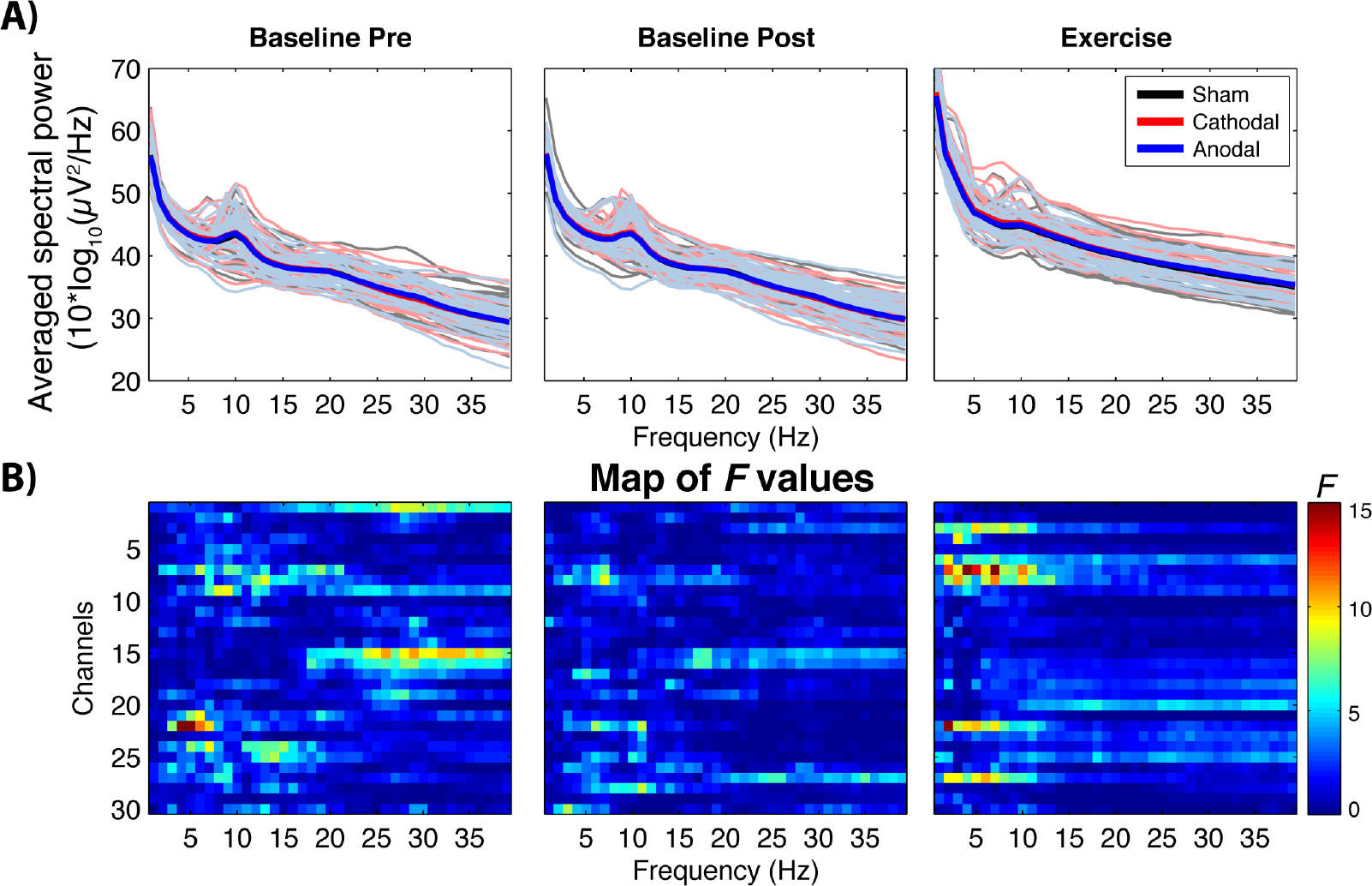
Differences in brain power spectrum as a function of tDCS condition. A) Average EEG power spectrum across participants among anodal (blue lines), cathodal (red line) and sham (black lines) condition at baseline pre, baseline post and exercise period. The shaded lines denote the average tonic spectral power for each participant and condition (given that there were not significant differences between conditions, the lines tend to overlap). B) Parametric F-test colormap comparing the relative power across frequency (x-axes) and channels (y-axes). Note that the analysis of the other periods (warmup and recovery) did not yield significant between-intensity differences.

The event-related spectral perturbation (stimulus-locked) analysis in the flanker task (see Fig. 3) did not reveal any main effect of condition for the congruent or incongruent trial (both ps > 0.05).

**Fig 3.**
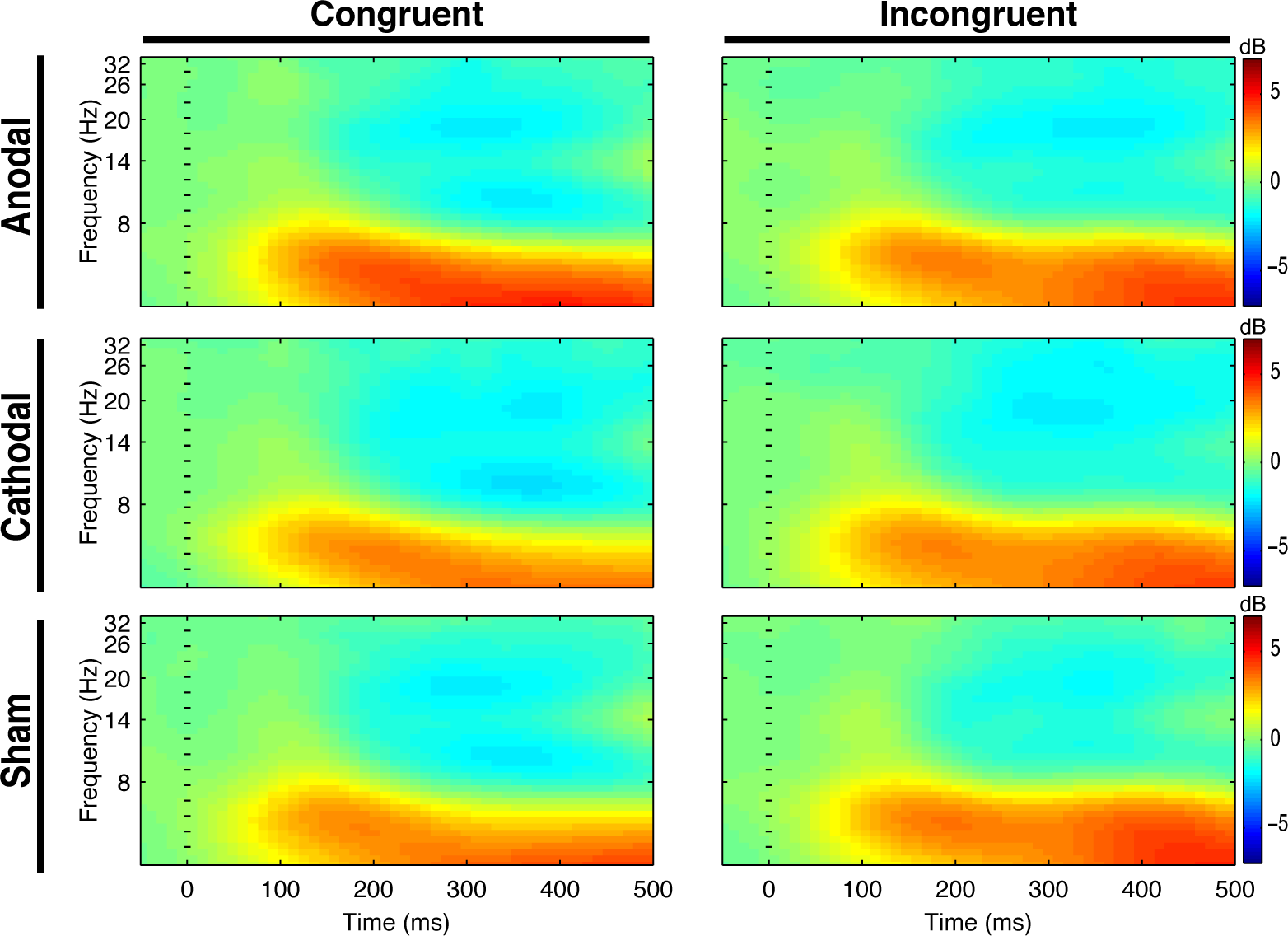
Event-related spectral perturbation during the flanker task. Time-locked spectral power averaged over all electrodes for each condition. Each panel illustrates time-frequency power across time (x-axes) and frequency (y-axes) for the congruent and incongruent stimuli (blue: decreases; red: increases). Dashed vertical line represents stimulus onset.

#### Executive task

A main effect of stimulus was reported in the flanker task, with participants being less accurate (M= 98 vs 91 % correct responses; F(2,34) = 13.17, p < 0.01) and slower (423 vs 515 ms; F(2,34) = 182.39, p < 0.05) in the incongruent stimulus compared to the congruent stimulus. There were no significant differences between conditions for the congruent and incongruent target, for RT and accuracy (Fs < 1, all ps > 0.05).

## Discussion

To the best of our knowledge, this is the first study testing the influence of prefrontal cortex tDCS’ stimulation on self-paced exercise and brain activity during exercise. The main finding of this study was that 20-min anodal or cathodal tDCS’ stimulation (relative to sham) over the left dorsolateral prefrontal cortex did not affect exercise performance or brain electrical activity. Moreover, neither sRPE, EEG or cognitive performance were affected by the stimulation. Our findings indicated that anodal or cathodal tDCS applied over the left dorsolateral prefrontal cortex before exercise did not modulate exercise performance during a 20-min time-trial self-paced exercise. This finding contrasts the results of the only previous study testing the effect of tDCS on cycling over the same brain area [13], as well as previous studies reporting positive effects of tDCS.

The rationale of our study was that the prefrontal cortex is involved in the control of self-paced exercise, and therefore stimulating it via tDCS would increase performance. In view of our null results, it may be possible that, through experience, self-pacing the effort became a more automatic task for our experienced cyclists, requiring less involvement of brain areas typically linked to executive processing. This may account for the apparent discrepancy of our results with those of the only previous study [13] testing the effect of tDCS over the prefrontal cortex on performance in a cycling task (a time to exhaustion test performed in the cycle ergometer). Indeed, participants in Lattari et al.’s study only reported 3 hours per week of aerobic physical activity the last six months, and hence could clearly not be classified as experienced cyclists. Therefore, the stimulation of the prefrontal cortex, instead of M1 as in the majority of previous positive findings, would explain the lack of effect of tDCS in our experiment.

Another factor that could help explaining our null results refers to the intensity of the stimulation. It is possible that 2mA (the most commonly used tDCS intensity in this research domain) was not high enough to affect neuronal circuits and hence to modulate exercise performance. Indeed, a study by Vöröslakos et al. [30] suggests that much higher current intensities are necessary to induce observable effects of electric brain stimulation. However, Vöröslakos et al. used transcranial alternating current stimulation (tACS) in their experiment which somewhat limits a direct comparison with studies using tDCS. It could be also possible that an individualized current intensity would be necessary to affect exercise performance due to the high inter-variability across participants (see [31], for discussion on this issue).

The hypothesis that anodal stimulation would increase EEG amplitude was not confirmed in the present study. After the 20-min stimulation, the EEG spectral power was similar across all condition for each period of time. This null effect is in line with the outcome of a review by Horvath et al. [34] who found that tDCS does not appear to modulate EEG power spectrum measures or event-related potential measures. This is also supported by the inconsistence aftereffect of tDCS on brain oscillations reported across studies [32]. Once again, the null effect of tDCS on the EEG signal could be explained by the low intensity of the stimulation. Indeed, using tACS, Vöröslakos et al. [30] found that currents between 4-6 mA should be delivered to modulate EEG amplitude.

The rationale of including the flanker task after the cycling self-paced exercise was that any change in physical performance and brain activity via tDCS would modulate the subsequent influence of cycling on cognitive (inhibition) performance. The lack of differences in physical exertion, RPE and EEG between the three experimental conditions make reasonable to have found no difference in RT or accuracy as a function of tDCS.

Apart from the abovementioned alternative explanations, we believe that a key methodological aspect could explain the discrepancy between our null findings and previous published studies, as well as the inconsistencies found in this literature (see [8] for discussion on this issue): the sample size of previous reports. The sample size of the vast majority of the tDCS studies in the Sport Science domain are low. According to a recent review, to date, the average sample size in tDCS’ experiment is N=14 [8]. If one assume that there is a true effect of tDCS over exercise performance, by testing 14 participants one would be assuming an effect size of dz=0.81 for a paired-sample two-tailed t-test (anodal vs. sham) and an a priori power of 1-β=.8 [18]. Testing a lower sample size (like in [9,11,13]) would assume an even larger effect size. However, such large effects are very unlikely in the tDCS research domain. For instance, the estimate average effect size for tDCS studies in cognition is dz=0.45 [33]. This would suggest, together with the low reproducibility of tDCS’ studies [33,34], that a statistically significant effect from a published tDCS-exercise study with a small sample size (which would not ensure sufficient statistical power) may easily reflect a false positive [35]. In view of our null result, one might wonder whether our study, assuming there is a true small effect of tDCS over self-paced exercise, was also underpowered even if we performed an a priori power analysis (based on an expected medium effect size). In that respect, it is worth noting that, to the best of our knowledge, our study has tested the largest sample size ever in this research domain. At this point, we believe that a meta-analytical review is necessary to unveil the overall effect (if any) of tDCS over exercise/sport performance and the effect of potential moderators (e.g., electrode site). Finally, we believe that the “file drawer effect” (i.e., the tendency to only publish positive outcomes) might be biasing the literature to positive findings [36].

## Conclusions

tDCS is an increasingly popular technique used within a wide range of settings, from treatment of neurological disorders, to attempting to improve exercise performance. Our data, however, add further to the mixed evidence in this area, challenging the idea that an acute session of tDCS can improve physical performance. At this point, we believe that research on this topic will benefit from further methodologically sound research in order to accumulate evidence on whether an acute session of tDCS affect sport performance or not.

## Practical applications

The use of tDCS is increasing in popularity in sport science

tDCS over the left prefrontal cortex does not improve performance in trained cyclists

tDCS does not seem to change EEG activity at rest or during exercise

## Acknowledgments

The authors thank Newronika S.L. for lending their tDCS device. The company was not involved in the study.

## Additional information

### Data availability

All data can be found in https://doi.org/10.5281/zenodo.1313703

